# Brain-derived neurotrophic factor supports pericyte and vascular homeostasis in the aging brain

**DOI:** 10.1101/2025.10.03.680251

**Authors:** Qinghua Luo, Wenqiang Quan, Qian Cao, Chris Scheffel, Wenlin Hao, Tomomi Furihata, Guoping Peng, Zhenyu Tang, Yang Liu

**Affiliations:** Department of Neurology, The second affiliated hospital of Nanchang University, Nanchang, China; Department of Neurology, Saarland University, Homburg, Germany; Department of Clinical Laboratory, Tongji Hospital, Tongji University Medical School, Shanghai, China; Center for Gender-Specific Biology and Medicine (CGBM), Saarland University, Homburg, Germany; Department of Clinical Pharmacy and Experimental Therapeutics, School of Pharmacy, Tokyo University of Pharmacy and Life Sciences, Tokyo, Japan; Department of Neurology, First Affiliated Hospital, Zhejiang University School of Medicine, Hangzhou, China

**Keywords:** Aging, BDNF, microcirculation, and pericyte

## Abstract

Microvascular circulation in the brain is often impaired in connection with the loss of pericytes in old age. The neurotrophic factor BDNF also decreases in the aging brain. We hypothesized that BDNF regulates the homeostasis of cerebral pericytes and microvasculature. We used differently aged C57BL/6J mice, and C57BL6 mice with conditional knockout of *Bdnf* gene. Collagen IV-positive microvessels and PDGFRβ-positive pericytes in the brain were counted after immunological staining. Pericytes were also quantified by Western blot of PDGFRβ and CD13 in isolated cerebral microvessels. The level of BDNF and TrkB phosphorylation was determined in brain homogenates. To demonstrate the direct effect of BDNF on pericytes, TrkB and pericytes were co-stained in brain tissue, single-cell sequencing and transcriptomic analysis were used to identify and characterize *Ntrk2*-expressing pericytes, and TrkB was also detected in the pericyte cell line by Western blot. Cultured pericytes were further treated with recombinant BDNF in the presence and absence of an Akt inhibitor and examined for PDGFRβ expression. The length and branching of microvessels and pericytes decreased in conjunction with the reduction in mature BDNF and TrkB phosphorylation in aging brains. Deficiency of BDNF in neurons or astrocytes was sufficient to reduce cerebral microvessels, PDGFRβ and CD13 concentrations and Akt and Erk1/2 phosphorylation in isolated blood vessels. A subset of pericytes in the brain and cultured pericytes expressed TrkB. BDNF treatment increased PDGFRβ expression along with Akt and Erk1/2 phosphorylation in cultured cells. The effect of BDNF on PDGFRβ expression was abolished by treatment with Akt inhibitor. Therefore, BDNF induces the expression of PDGFRβ and CD13 by activating Akt signaling in pericytes, promoting the homeostasis of pericytes and microvasculature in the aging brain. Our study identified a BDNF-mediated mechanism that regulates microvascular integrity in the aged brain.

## Introduction

Aging is the non-modifiable risk factor for both large artery disease [1], and small vessel disease, which often shows up as white matter hyperintensities (WMH) on brain imaging [2]. Longitudinal studies have indicated that increasing WMH volume predicts cognitive decline, stroke and death [3, 4]. Aging is also the risk factor for Alzheimer’s disease (AD) [5]. Notably, many AD patients exhibit WMH on brain imaging [6, 7]. Although the biological basis of WMH remains unclear, it is thought that ischemia or hypoperfusion, causing demyelination and axonal loss, and disruption of blood-brain barrier (BBB) are the main pathological changes. The pathologies of capillaries, especially endothelial cells and pericytes, should be considered [8, 9]. The cerebrovascular impairment also applies to mice during aging. For example, both vascular length and branching density in various brain regions decrease by ∼10 % in 18-month-old mice compared to 2-month-old controls [10]. Our previous studies have also shown that blood perfusion and vasculature in the brain are reduced in both APP/PS1 and tau-transgenic AD mice [11]. Therefore, mechanisms that promote vascular health in the brain during aging should be explored.

Pericytes surrounding the endothelial cells of the brain capillaries are essential for the structural and functional integrity of the microcirculation in the brain and thus contribute to the health of the vascular system [12]. Pericytes express platelet-derived growth factor receptor β (PDGFRβ), which releases into the cerebrospinal fluid (CSF) when cells are injured. Indeed, soluble PDGFRβ in CSF increases during aging in association with neuroinflammatory activation and BBB damage [13]. Soluble PDGFRβ also increases in CSF at a very early stage in old adults with cognitive deficits [14]. Similarly, the density of pericytes decreases in the deep cortical layers, hippocampal network and basal forebrain areas in aged mouse brains, where blood extravasation is increased and the baseline and on-demand blood oxygenation are reduced [10]. Therefore, we decided to explore molecular mechanisms that regulate the healthy activity of pericytes during aging.

Brain-derived neurotrophic factor (BDNF) is expressed by neurons and glial cells (e.g., astrocytes and microglia) and plays an important role in neuronal development, differentiation, maintenance and plasticity throughout life [15]. BDNF is produced as a precursor protein (pro-BDNF) and cleaved to yield the mature isoform (mBDNF). Binding of mBDNF to tyrosine kinase B receptor (TrkB) induces the phosphorylation of intracellular tyrosine residues and the subsequent downstream kinases such as PI3K/Akt, whose activation exerts anti-apoptotic and pro-survival effects [16]. Interestingly, BDNF also promotes survival and proliferation of vascular endothelial and smooth muscle cells in culture [17, 18]. Knockout of *Bdnf* gene leads to endothelial cell apoptosis and reduces endothelial cell-cell contacts in the intramyocardial arterioles and capillaries of mice during late embryogenesis, resulting in hemorrhage in the ventricular wall, decreased cardiac contractility, and early postnatal death [17]. Since *BDNF* transcription was substantially downregulated in various cortical regions (e.g., entorhinal cortex, superior frontal gyrus, and postcentral gyrus), and the hippocampus in aged individuals compared to young individuals [19], we hypothesized that the reduction in BDNF may also affect pericytes and vasculature in the brain during aging. We did not exclude the important effect of BDNF on endothelial cells in aging-related cerebrovascular changes; however, we focused on pericytes in this project.

We determined the amount of vasculature and PDGFRβ and CD13 expression in pericytes in correlation with BDNF protein levels in mouse brains of different ages. We then analyzed the effects of BDNF deficiency on the cerebral vasculature and pericyte density. Finally, we validated the effect of BDNF on PDGFRβ expression in cultured human pericytes. Our project demonstrated that BDNF favors vascular and pericyte health in the aged brain.

## Materials and Methods

### Animal models and cross-breeding

C57BL6/J mice for vascular analysis at different ages were originally ordered from Charles River Laboratories (Sulzfeld, Germany). *Bdnf*^fl/fl^ mice carrying loxP-site-flanked exon IX, the single protein coding exon of *Bdnf* gene, were kindly provided by M. Sendtner (University of Würzburg) [20]. *Camk2a*-CreERT2 transgenic mice, expressing a fusion protein of Cre recombinase and an estrogen receptor ligand binding domain (CreERT2) under the control of the mouse *Calcium/calmodulin-dependent protein kinase II α* promoter, were obtained from the Jackson Laboratory (Bar Harbor, USA; Stock Number: 012362) [21]. *GFAP*-CreERT2 transgenic mice, expressing CreERT2 under the control of human *GFAP* promoter, were kindly provided by F. Kirchhoff, Saarland University [22]. The cross-breeding between *Bdnf*^fl/fl^ and *Camk2a*-CreERT2 or *GFAP*-CreERT2 have been conducted in our previous study [23]. Mice expressing CreERT2 and their matched control littermates without Cre expression were injected with tamoxifen (Sigma-Aldrich Chemie GmbH, Munich, Germany; 100 mg/kg) in corn oil once daily for 5 days at 7 months of age and analyzed for phenotype when they were 10 months old.

To detect TrkB expression on pericytes, Ai14 Cre reporter mice were mated with PDGFRβ-P2A-CreERT2 mice to obtain tdTomato^fl/fl^Cre^+/-^ genotype (both mice were imported from the Jackson Laboratory, Bar Harbor, ME, USA, with the strain numbers: 007914 and 030201, respectively) [21, 24]. After injection of tamoxifen as described above, tdTomato^fl/fl^Cre^+/-^ mice expressed tdTomato specifically in pericytes.

Animal breeding, experimental procedure and methods of killing were conducted in accordance with national rules and ARRIVE guidelines, and were authorized by Landesamt für Verbraucherschutz, Saarland, Germany (registration numbers: 06/2017, and 05/2022) and the ethical committee in Nanchang University, China.

### Tissue collection and isolation of blood vessels

Mice were euthanized by inhalation of overdose isoflurane and perfused with ice-cold phosphate-buffered saline (PBS). The brain was removed and divided. The left hemisphere was immediately fixed in 4% paraformaldehyde (Sigma-Aldrich Chemie GmbH) in PBS and embedded in paraffin or Tissue-Tek® O.C.T. Compound (Sakura Finetek Europe B.V., AJ Alphen aan den Rijn, the Netherlands) for histological analysis. The right hemisphere was snap-frozen in liquid nitrogen and stored at -80°C until biochemical analysis.

To isolate microvessels from the brain, the cortex and hippocampus from right hemisphere were carefully dissected and brain vessel fragments were isolated using our established protocol [25]. Briefly, brain tissues were homogenized in HEPES-contained Hanks’ balanced salt solution (HBSS) and centrifuged at 4,400 g in HEPES-HBSS buffer supplemented with dextran from *Leuconostoc spp.* (molecular weight ∼ 70,000; Sigma-Aldrich Chemie GmbH) to delete myelin. Vessel fragments were re-suspended in HEPES-HBSS buffer supplemented with 0.1% bovine serum albumin (Sigma-Aldrich Chemie GmbH) and filtered by nylon mesh filter. The filtrates passing through 100 but not 20 μm-meshes were collected and stored at -80°C for biochemical analysis.

### Histological analysis

To quantify vasculature in the brain, our established protocol was used [26, 27]. Thirty-μm-thick sagittal sections were serially cut from the paraffin-embedded left hemisphere. Four serial sections per mouse with 300µm of distance in between were deparaffinized, heated at 80°C in citrate buffer (10mM, pH = 6) for 1 hour and digested with Digest-All 3 (Pepsin) (Thermo Fisher Scientific) for 20 minutes. After blocking with 0.2% casein in PBS/0.3% Triton X-100, brain sections were stained with rabbit anti-collagen IV polyclonal antibody (Catalog number: ab6586; Abcam, Cambridge, UK), biotin-conjugated goat anti-rabbit IgG and Cy3-conjugated streptavidin (both from Jackson ImmunoResearch Europe Ltd, Cambridgeshire, UK). After mounting, the entire hippocampus was imaged with Microlucida on a Zeiss AxioImager.Z2 microscope equipped with a Stereo Investigator system (MBF Bioscience, Williston, VT, USA). Blood vessels larger than 6µm in diameter were cut away before subsequent quantification. The length and branch points of the collagen type IV-positive blood vessels were analyzed using a free software, AngioTool (http://angiotool.nci.nih.gov) [28]. The parameters of analysis for all compared samples were kept constant. The length and branching points were adjusted by area of interest.

To determine the density of pericytes on blood vessels, twenty-μm-thick sagittal sections were serially cut from the frozen left hemisphere. Three serial sections per mouse with 300µm of distance in between were heated at 80°C in citrate buffer (10mM, pH = 6) for 30 minutes. After blocking with 0.2% casein in PBS/0.3% Triton X-100, brain sections were serially stained with rabbit anti-PDGFRβ monoclonal antibody (clone 28E1; Cell Signaling Technology Europe B.V., Leiden, The Netherlands) and anti-CD31 monoclonal antibody (clone D8V9E; Cell Signaling Technology) with relevant fluorescence-conjugated second antibodies. Ten areas in the hippocampus were randomly chosen and imaged under a 40× objective with the Stereo Investigator system. To better present the images on the fluorescent staining, stack images were acquired with an interval of 2 µm for 5 layers, deconvoluted and Z-projected with maximum intensity. The total length of CD31-stained vessels was measured and PDGFRβ-positive cells on the vessels were counted.

To detect TrkB expression on pericytes, 30-μm-thick sagittal sections was cut from paraffin-embedded brain tissues of tamoxifen-injected tdTomato^fl/fl^Cre^+/-^ mice. After deparaffinization and antigen retrieval, brain sections were incubated with mouse monoclonal antibody against TrkB (clone 75133; Bio-Techne GmbH, Wiesbaden, Germany), and then biotin-conjugated goat anti-mouse IgG and Cy3-conjugated streptavidin (both from Jackson ImmunoResearch). Thereafter, the brain section was further stained with rabbit anti-red fluorescence protein (Catalog number: 600-401-379, Rockland Immunochemicals, Pottstown, USA) and Alexa488-conjugated goat anti-rabbit IgG (Jackson ImmunoResearch). The expression of TrkB on pericytes was presented by the colocalization of red and green fluorescence.

### Western blot analysis

Frozen brain tissues were homogenized on ice in radioimmunoprecipitation assay (RIPA) buffer supplemented with protease inhibitor cocktail and phosphatase inhibitors (50nM okadaic acid, 5mM sodium pyrophosphate, and 50mM NaF; Sigma-Aldrich Chemie GmbH), followed by centrifugation at 16,000 × g for 30 minutes at 4 °C to collect the supernatants. Isolated blood vessels were directly lysed in 2 × SDS-PAGE sample loading buffer containing 4% SDS and sonicated before loading. The protein level of BDNF was detected with rabbit polyclonal antibodies (Catalog number: NBP1-46750; Bio-Techne GmbH). Phosphorylated TrkB (pTyr817) and total TrkB were detected with rabbit monoclonal antibodies: clone SC0556 (Bio-Techne GmbH) and clone 80E3 (Cell Signaling Technology), respectively. For the detection of proteins in cerebral capillaries, rabbit monoclonal antibodies against PDGFRβ, CD13/APN, phosphorylated Akt (Ser473), and phosphorylated Erk1/2 (Thr202/Tyr204) (clones 28E1, D6V1W, D9E, and D13.14.4E, respectively; Cell Signaling Technology), rabbit polyclonal antibody against Akt (Catalog number: 9272; Cell Signaling Technology), and mouse monoclonal antibody against Erk1/2 (clone L34F12; Cell Signaling Technology) were used. For loading controls, rabbit monoclonal antibodies against GAPDH and β-actin (clones 14C10 and 13E5, respectively; Cell Signaling Technology) and mouse monoclonal antibody against α-tubulin (clone DM1A; Abcam) were used. Western blots were visualized via the ECL method (PerkinElmer LAS GmbH, Rodgau, Germany). Densitometric analysis of bands was performed with Image J software. For each sample, the protein level was calculated as a ratio of target protein/β-actin, α-tubulin or GAPDH.

### Single-cell sequencing and transcriptomic analysis of pericytes and endothelial cells

The single-cell RNA-seq dataset was downloaded from the Neuroscience Multi-omic Data Archive (https://assets.nemoarchive.org/dat-61kfys3) [29]. The experimental procedures and initial computational analysis, including 10× Genomics library preparation, sequencing, quality control, and normalization, are detailed in the original publication’s methods section [29]. The dataset comprised 1.2 million high-quality cells from young (2-month) and aged (18-month) mouse brains. Raw UMI counts were normalized using Counts Per Million followed by log2 transformation to account for sequencing depth variation. Cell annotations were assigned based on the Allen Brain Cell - Whole Mouse Brain Atlas reference.

From the annotated dataset, we isolated 17,187 pericytes (young: 5,813; aged: 11,374) and 51,454 high-quality endothelial cells (young: 22,898; old: 28,556), classified under the vascular cell hierarchy (vascular → pericytes and vascular → endothelial cells) using Scanpy 1.9.8 (Python 3.8.20). To ensure data robustness, a multi-tiered quality control protocol was implemented: (1) initial filtration retained only cells with > 500 detected genes, > 1,000 UMIs, and < 10% mitochondrial gene content; (2) two independent Scrublet analyses consistently demonstrated 0% doublet rates; and (3) cell identities were validated by plotting the expression of key marker genes for pericytes (*Pdgfrb*, *Rgs5*, and *Des*) and endothelial cells (*Cldn5*, *Flt1*, and *Pecam1*) on the UMAP coordinates.

Comparative transcriptomic analysis between age groups was performed using a standardized analytical pipeline. The different transcription of *Ntrk2* and *Ngfr* genes, encoding receptors for BDNF was also analyzed. Differential expression testing was conducted using the Wilcoxon rank-sum test (implemented in Scanpy), with a pre-filtering step to exclude genes detected in **<** 10% of cells. Significant differentially expressed genes (DEGs) were defined as those with |log2(fold change)| > 1 and Benjamini-Hochberg-adjusted *p* < 0.05. The non-parametric approach was chosen for its robustness to zero-inflated single-cell data distributions [30].

DEGs were further subjected to functional enrichment. Gene Ontology (GO) and Kyoto Encyclopedia of Genes and Genomes (KEGG) pathway enrichment analyses were performed using the R package clusterProfiler [31]. Gene annotation was based on the org.Mm.eg.db database (Bioconductor, version 3.19.1), with the organism code mmu (Mus musculus) applied for KEGG enrichment. All analyses were conducted in R (version 4.4.0) within the Bioconductor framework (version 3.19). Significant pathways were identified through hypergeometric testing (*p* < 0.05), with pathway size filtering (5 - 500 genes) to exclude overly broad or narrow categories. Results were visualized as bar plots sorted by enrichment score (- log10(*p*-value)), annotated with gene ratios (observed/expected) and Benjamini-Hochberg-adjusted *p*-values. To ensure reproducibility, all analysis and visualization code - including volcano plots highlighting top DEGs - was implemented using ggplot2 (version 3.4.4).

The workflow of scRNA-seq data analysis is presented in Supplementary Figure 1.

### Pericyte culture and treatments

Human primary brain vascular pericytes (HBPC) were immortalized by infecting cells with tsSV40T lentiviral particles [32]. The selected immortalized HBPC clone 37 (hereafter referred to as HBPC/ci37) was used for our study. HBPC/ci37 cells were cultured at 33 °C with 5% CO2/ 95% air in pericyte medium (Catalog number: 1201; Sciencell Research Laboratories, Carlsbad, CA, USA) containing 2% (v/v) fetal bovine serum, 1% (w/v) pericyte growth factors, and penicillin-streptomycin. Culture flasks and plates were coated with Collagen Coating Solution (Catalog number: 125-50; Sigma-Aldrich). HBPC/ci37 cells were used at 40 ∼ 60 passages in this study.

To investigate the effects of BDNF stimulation on expression of PDGFRβ, pericytes were cultured in 12-well plate at 5.0 × 10^5^ cells/well. Before experiments, the culture medium was replaced with serum-free pericyte medium and cells were cultured at 37°C for 3 days to facilitate the cell differentiation [32]. Thereafter, pericytes were treated with recombinant human BDNF (Catalog number: 248-BDB-010; R&D Systems, Wiesbaden, Germany) at 0, 10, 50 and 100 ng/ml for 24 hours. At the end of experiments, cultured cells were lysed in RIPA buffer supplemented with protease and phosphatase inhibitor cocktail. Protein levels of TrkB, PDGFRβ, phosphor-/total-Akt and phosphor-/total-Erk1/2 were detected with quantitative Western blot as described in section “**Western blot analysis**”.

To investigate the effect of Akt activation on BDNF-induced PDGFRβ expression, cultured pericytes in 12-well plates were treated with BDNF at different concentrations with and without the presence of 1 μM AKT inhibitor VIII (Catalog number: 124018; Sigma-Aldrich Chemie GmbH) for 24 hours. Thereafter, cells were lysed for Western blot analysis.

### Statistical analysis

Data were presented as mean ± SEM. For multiple comparisons, we used one-way ANOVA followed by Bonferroni, Tukey, or Dunnett T3 post-*hoc* test (dependent on the result of Levene’s test to determine the equality of variances). Two independent-samples were compared with Students *t*-test or Mann-Whitney U test, depending on the distribution of values. All statistical analyses were performed with GraphPad Prism 8 version 8.0.2. for Windows (GraphPad Software, San Diego, CA, USA). Statistical significance was set at *p* < 0.05.

## Results

### Aging reduces vasculature and pericyte expression of PDGFRβ in the brain

To investigate the effects of aging on the cerebral vasculature, we first stained brain tissue from 6, 12 and 24 month old C57BL6/J mice for collagen type IV (Fig. 1, A). The length and density of branch points, adjusted for the region of interest, were similar in 6- and 12-month-old mice (6 vs. 12 months: length, 7.40 ± 0.29 vs. 7.49 ± 0.22 arbitrary unit [A.U.], *t*-test, *p* = 0.846; branch points, 2.43 ± 0.31 vs. 2.15 ± 0.20 [A.U.], *t*-test, *p* = 0.521). To reduce the number of experimental animals, we combined the results for these two groups of young mice. We found that both the length and density of branch points of microvessels in the hippocampus of 24-month-old mice were significantly reduced compared to the 6-12-month-old control animals (Fig. 1, B and C; *t*-test, *p* < 0.05). Our findings are consistent with a recent study in which cerebral vasculature and branch point density were ∼10% lower in 18-month-old mice than in 2-month-old control mice, as determined by serial two-photon tomography imaging after cerebral vessels were labelled by cardiac perfusion of fluorescein isothiocyanate-conjugated albumin gel [10].

**Figure 1.**
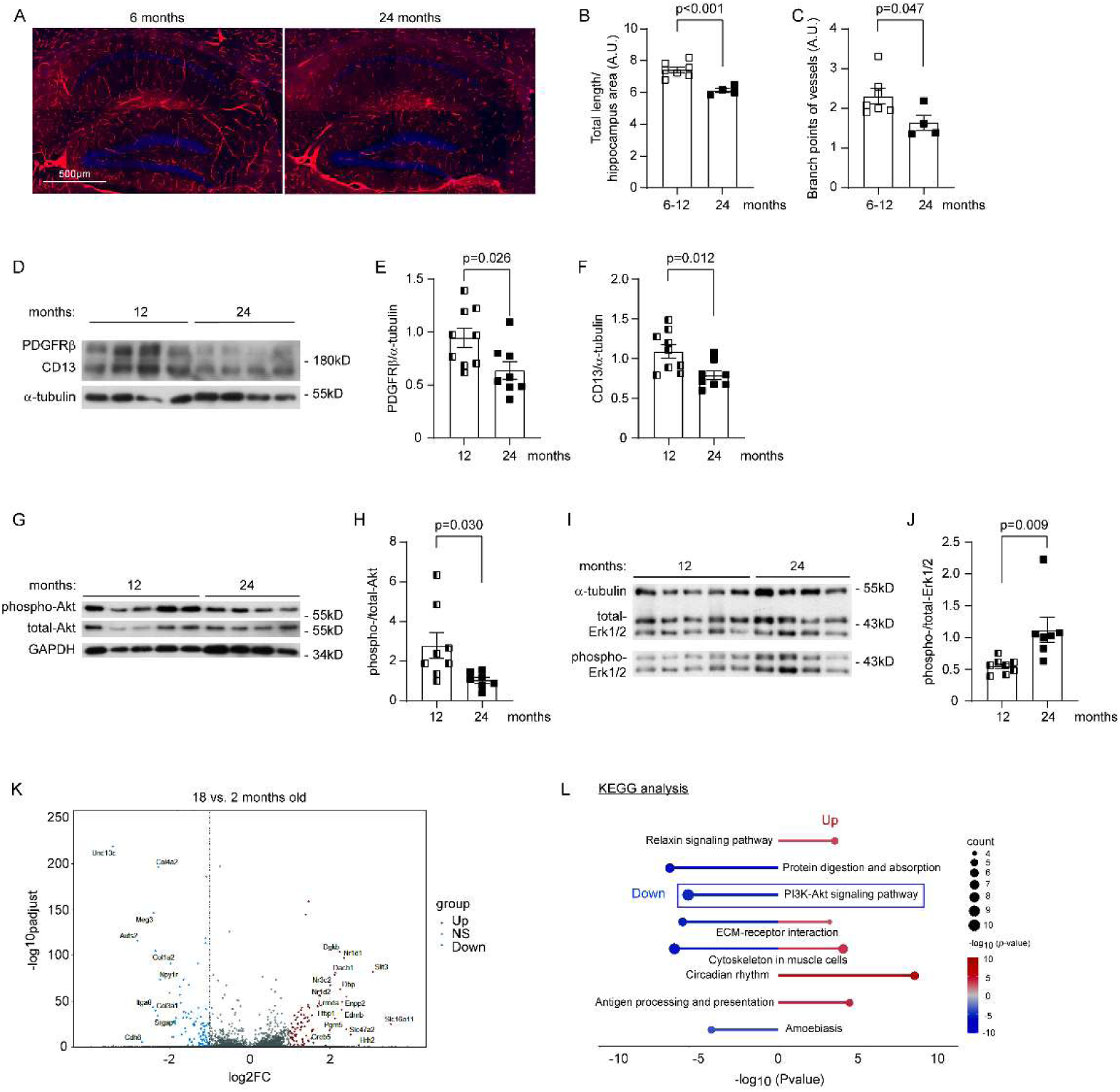
Aging reduces vasculature and pericyte expression of PDGFRβ in the brain. Brain tissue from C57BL6/J mice at 6, 12 and 24 months of age was stained for collagen type IV (A) and quantified for total length and branch points of microvessels (B and C; *t*-test, *n* = 7 and 4 for the 6-12 months and 24 months groups, respectively). Cerebral microvessels were isolated from 12 and 24-month-old mice and homogenized for Western blot analysis of pericyte markers PDGFRβ and CD13, and protein levels of phosphorylated and total Akt and Erk1/2 (D - J; *t*-test, *n* = 8 - 9 and 7 - 8 per group for the 6-12 months and 24 months groups, respectively). In following experiments, transcriptomic analysis was performed using a single-cell sequencing dataset. Volcano plots show up- and down-regulated DEGs in the brains of 18-month-old mice compared with 2-month-old mice (K; NS, genes with non-significant changes). The bar chart displays the top five significantly enriched KEGG pathways ranked by -log10 (*p*-value) (bar length). The size of adjacent circles corresponds to the number of DEGs in each pathway. KEGG pathway analysis indicated that the down-regulated genes are associated with PI3K-AKT signaling pathway (L).

Since pericytes regulate the structure and function of brain capillaries [12], we detected pericyte markers PDGFRβ and CD13 in the cerebral microvessels isolated from 12 and 24-month-old C57BL6/J mice by Western blot. We observed that the protein levels of both PDGFRβ and CD13 were significantly lower in 24-month-old mice than in 12-month-old control animals (Fig. 1, D - F; *t*-test, *p* < 0.05). In further experiments, we determined that aging inhibited the activation of Akt but enhanced the activation of Erk1/2, as the phosphorylation level of Akt decreased in the cerebral microvessels of 24-month-old C57BL6/J mice compared with 12-month-old control mice, but that of Erk1/2 increased (Fig. 1, G - J; *t*-test, *p* < 0.05), indicating a possible role of Akt inhibition in the reduction of vessels in the aged brain.

We performed a further transcriptomic analysis of a single-cell sequencing dataset generated by another laboratory [29] to confirm our findings on the effects of ageing on pericytes. A total of 186 DEGs (99 downregulated and 87 upregulated) with |log2(fold change)| > 1 and Benjamini-Hochberg-adjusted *p* < 0.05 were identified in pericytes from the whole brain of 18-month-old C57BL6/J mice compared with 2-month-old control mice (Fig. 1, K). KEGG pathway analysis was conducted separately for downregulated and upregulated DEGs. KEGG analysis showed that both up- and down-regulated DEGs were involved in the “extracellular matrix-receptor interaction” and “cytoskeleton in muscle cells”, suggesting effects of aging on pericyte signaling and contraction. Of note, aging down-regulated the transcritption of *Col4a1*, *Col4a2*, *Col1a2*, *Itga8*, *Lamb1*, *Igf2*, *Magi1*, *Angpt1*, *Itga9*, and *Tek* genes associated with the PI3K-Akt signaling pathway (Fig. 1, L), consistent with the above observations in the quantitative Western blot analysis that aging reduced phosphorylation of Akt in the isolated microvessels (Fig. 1, G). Similarly, genes related to protein digestion and absorbation and amoebiosis were reduced, while relaxin signaling appeared to be activated (Fig. 1, L). Relaxin activates PI3K, which is involved in the induction of matrix metalloproteinases [33]. Relaxin treatment also enhances endothelium-dependent relaxation and decreases myogenic tone in resistance arteries [34], although the effects of relaxin on pericytes are unknown. We observed that aging upregulated the transcription of *Hspa2*, *H2-D1*, *B2m*, *H2-K1* and *Hspa1b* genes, which are related to antigen processing and presentation (Fig. 1, L). It remains to be investigated whether aging strengthens immune signaling pathway in pericytes.

### Aging reduces BDNF signaling in the brain

Since BDNF has the potential to regulate angiogenesis [17, 18] and BDNF expression is reduced by aging in the human brain [19], we were interested to examine the BDNF expression and signaling in brains from differently aged mice. By quantitative analysis of brain homogenates from 6, 12 and 24-month-old C57BL6/J mice with Western blot (Fig. 2, A), we observed that the protein levels of mature BDNF (mBDNF), but not BDNF precursor (pro-BDNF) decreased along with aging (Fig. 2, A - C; one-way ANOVA, *p* < 0.05). mBDNF binds to TrkB with a high affinity and induces phosphorylation of intracellular tyrosine residues of the receptor [16]. Indeed, the phosphorylation level of TrkB dramatically decreased in the brain homogenates of 12 and 24-month-old mice compared to the younger 6-month-old control animals and in an age-dependent manner (Fig. 2, A and D; one-way ANOVA, *p* < 0.05).

**Fig. 2.**
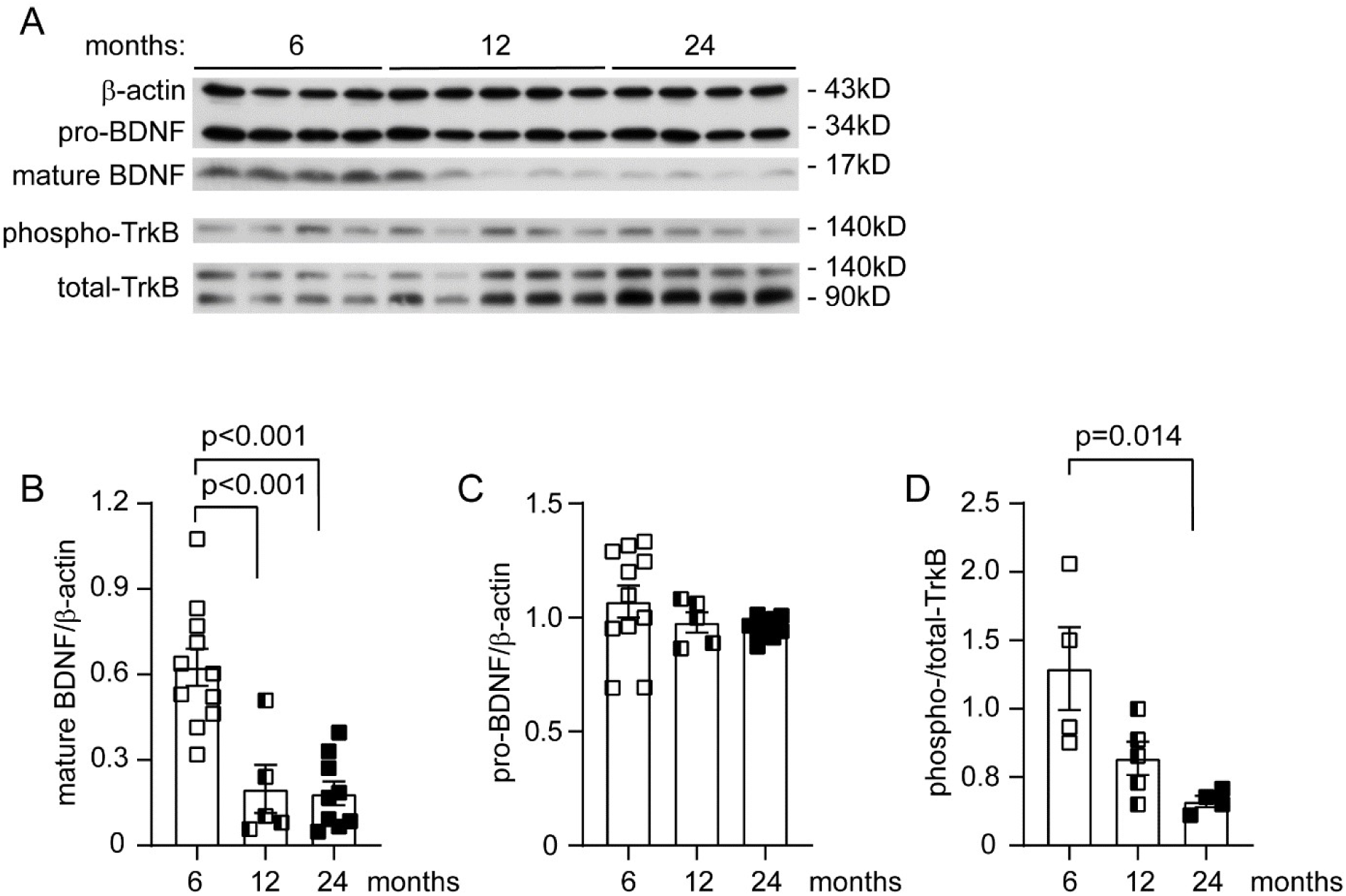
Aging reduces BDNF maturation and signaling in the brain. Brains of C57BL6/J mice at 6, 12 and 24 months of age were homogenized for quantitative Western blot of BDNF (A). Normal ageing significantly reduced the protein levels of maturated BDNF, but not pro-BDNF (B and C; one-way ANOVA followed by Bonferroni post-hoc test; *n* = 5 - 11 per group). Moreover, aging significantly decreased the phosphorylation level of TrkB, the specific receptor of BDNF (A and D; one-way ANOVA followed by Bonferroni post-hoc test; *n* = 4 - 5 per group).

### Deficiency of neuronal BDNF reduces vasculature and pericytes in the brain

After we had observed that aging reduced: 1) vasculature and pericytes, and 2) BDNF signaling in the brain, we asked whether BDNF regulated cerebral vasculature. We have established C57BL6 mice with Bdnf^fl/fl^/Camk2a-CreERT2^tg^ and Bdnf^fl/fl^/Camk2a-CreERT2^wt^ genotypes. Both groups of mice were injected with tamoxifen (*i.p.*) at 7 months of age to specifically delete BDNF in neurons, as we have shown in a previous study [23]. The mice were analyzed 3 months later (at 10 months of age). As shown in Fig. 3, A - C, deficiency of BDNF in neurons significantly reduced the length of microvessels (*t*-test, *p* < 0.05) and tended to decrease the density of branch points (*t*-test, *p* = 0.05) in the hippocampus. By counting PDGFRβ-positive cells on CD31-positive vessels, we also found that the lack of neuronal BDNF reduced the number of pericytes per unit length of blood vessels (Fig. 3, D and E; *t*-test, *p* < 0.05). In Western blot analysis of isolated brain microvessels, protein levels of both PDGFRβ and CD13 were lower in neuronal BDNF-deficient mice than in BDNF wild-type control mice (Fig. 3, F - H; *t*-test, *p* < 0.05), consistent with the pericyte count described above. Interestingly, deficiency of BDNF in neurons significantly reduced the phosphorylation of Akt, but not Erk1/2 in the isolated blood vessels as compared with neuronal BDNF wild-type mice (Fig. 3, F and I; *t*-test, *p* < 0.05).

**Figure 3.**
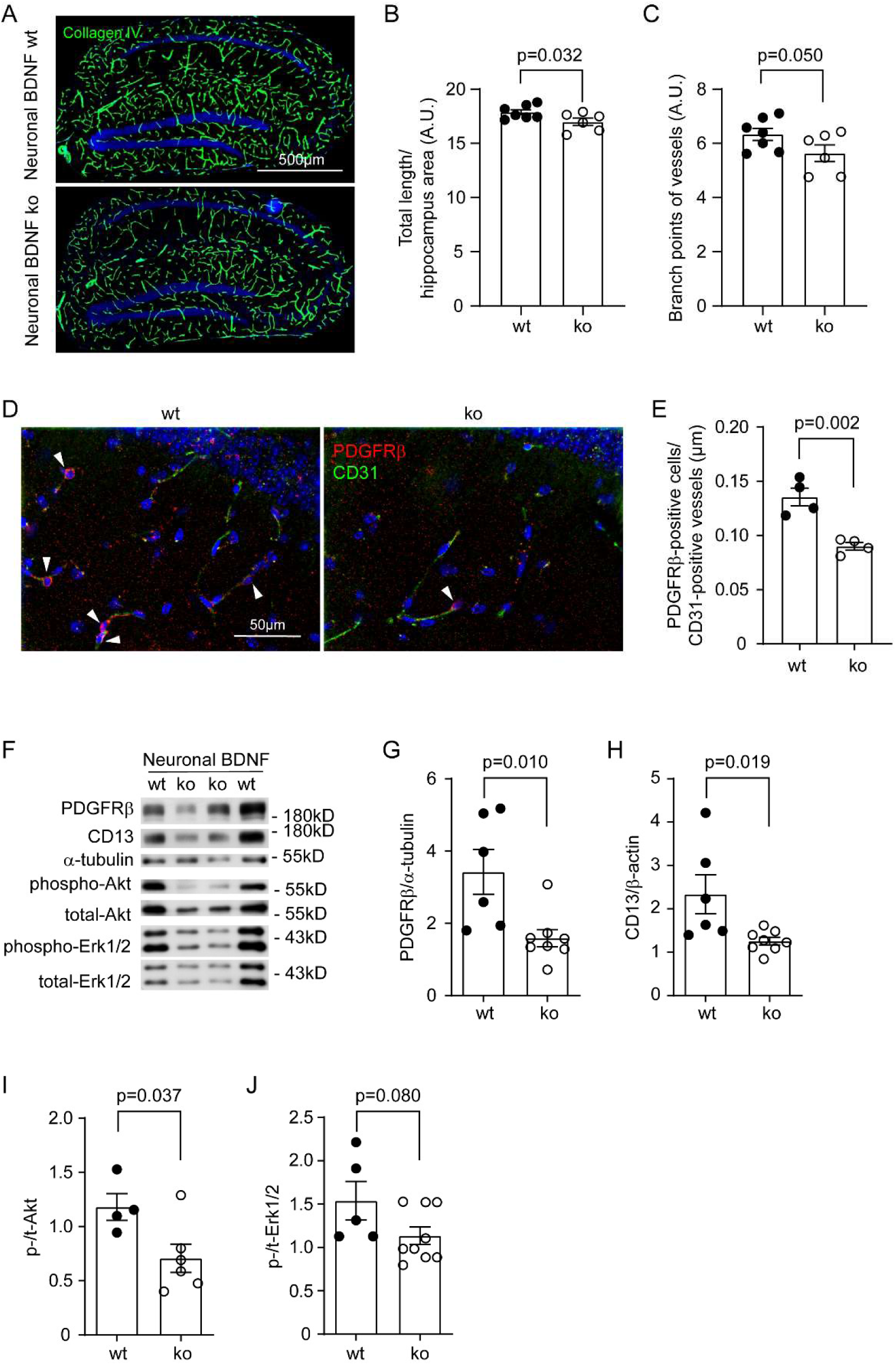
Deficiency of neuronal BDNF reduces vasculature and pericytes in the brain. Ten-month-old C57BL6 mice with (ko) and without (wt) knockout of *Bdnf* gene in neurons for 3 months were stained for collagen type IV and quantified for the vasculature (A). Deficiency of neuronal BDNF significantly reduced the total length of cerebral vessels and tended to decrease the density of branch points of vessels (B and C; *t*-test, *n* = 6 - 7 per group). The brain sections were also co-stained for PDGFRβ and CD31. PDGFRβ-positive pericytes were counted and adjusted by the length of CD31-positive vessels (D). Deficiency of neuronal BDNF significantly reduced the number of pericytes (E; *t*-test, *n* = 4 per group). Additionally, microvessels were isolated from brains and detected with Western blot for pericyte markers and relevant signaling molecules (F). Deficiency of neuronal BDNF decreased protein levels of PDGFRβ and CD13 (G and H; *t*-test, *n* = 6 - 8 per group), as well as the phosphorylation of Akt, but not Erk1/2 (I and J; *t*-test, *n* = 4 - 9 per group).

### Deficiency of astrocyte BDNF reduces vasculature and pericytes in the brain

In our previous study, we have observed that BDNF is also expressed in astrocytes [23]. Since astrocytes (particularly their endfeet) surround capillaries, we hypothesized that astrocyte BDNF could also affect the generation of cerebral vasculature. C57BL6 mice with Bdnf^fl/fl^/Gfap-CreERT2^tg^ and Bdnf^fl/fl^/Gfap-CreERT2^wt^ genotypes have been established in our previous study [23]. After intraperitoneal injection of tamoxifen at 7 months of age, we could clearly observe that the protein levels of mature BDNF but not pro-BDNF were reduced in Bdnf^fl/fl^/Gfap-CreERT2^tg^ mice compared with Bdnf^fl/fl^/Gfap-CreERT2^wt^ mice when they were 10 months old (Fig. 4, A - C; *t*-test, *p* < 0.05). After immunostaining of collagen type IV, we observed that both the length and density of branch points of the microvessels were significantly fewer in astrocyte BDNF-deficient mice than in the BDNF wild-type control mice (Fig. 4, D - F; *t*-test, *p* < 0.05). In quantitative Western blot analysis of isolated blood vessels, we found that deficiency of BDNF in astrocytes significantly decreased the protein level of PDGFRβ but not CD13 (Fig. 4, G - I; *t*-test, *p* < 0.05). Deficiency of BDNF in astrocytes also significantly reduced phosphorylation of both Akt and Erk1/2 in the cerebral blood vessels (Fig. 4, J - L; *t*-test, *p* < 0.05).

**Figure 4.**
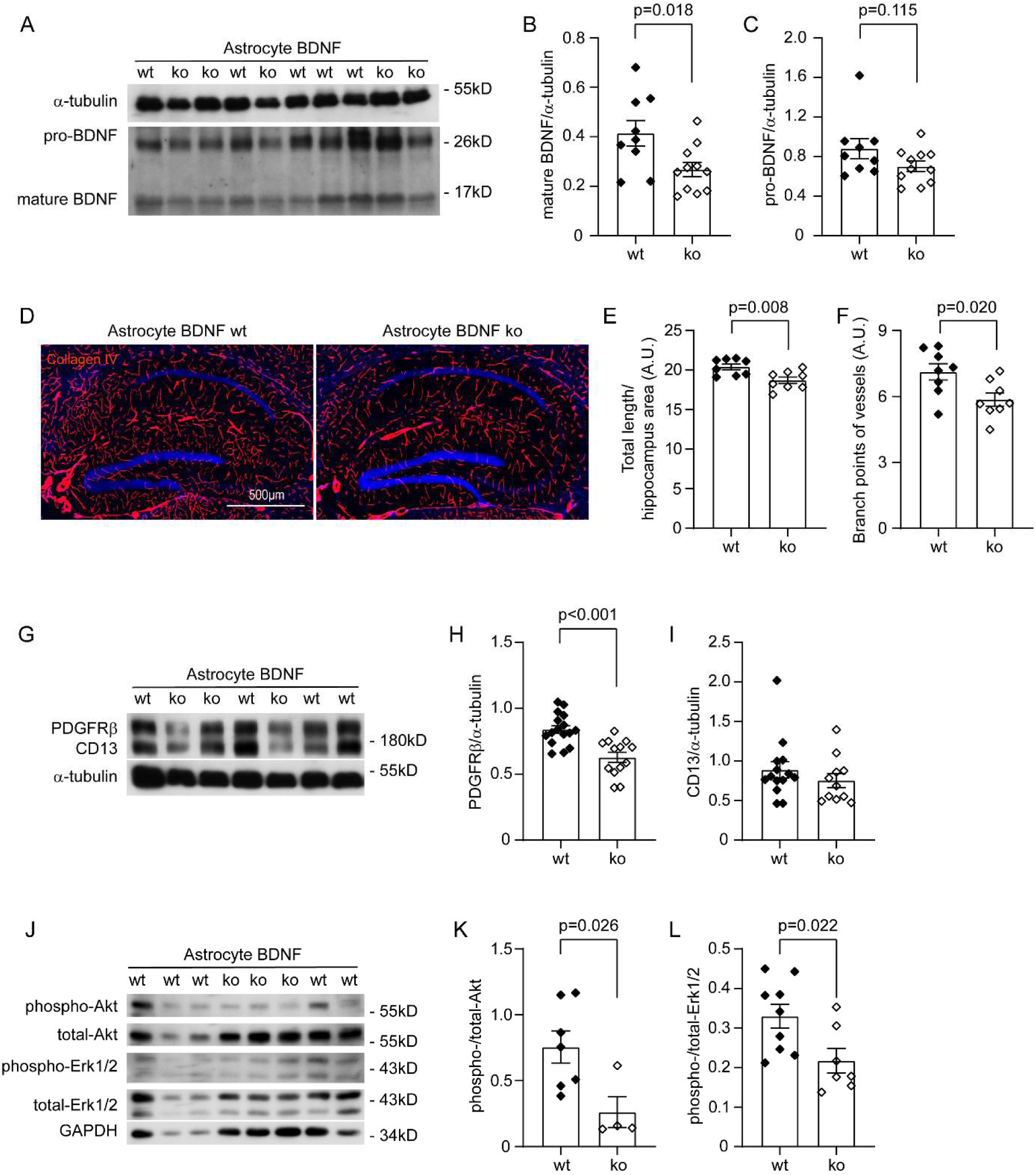
Deficiency of astrocyte BDNF reduces vasculature and pericytes in the brain. Brain homogenates from 10-month-old C57BL6 mice with (ko) and without (wt) knockout of *Bdnf* gene in astrocytes for 3 months were detected for protein levels of BDNF (A). Knockout of *Bdnf* gene significantly reduced mature BDNF but not pro-BDNF (B and C; *t*-test, *n* = 9 - 11 per group). Brain sections were then stained for collagen type IV and quantified for the vasculature (D). Deficiency of astrocyte BDNF significantly reduced both the length and density of branch points of cerebral vessels (E and F; *t*-test, *n* = 8 per group). Additionally, microvessels were isolated from brains and detected with Western blot for pericyte markers and relevant signaling molecules (G). Deficiency of astrocyte BDNF decreased the protein level of PDGFRβ, but not CD13 (H and I; *t*-test, *n* = 13 - 16 per group), and reduced the phosphorylation of both Akt and Erk1/2 (I and J; *t*-test, *n* = 4 - 9 per group).

### BDNF acts directly on pericytes

In aging and *Bdnf*-knockout animals, we have observed that BDNF correlated with the amount of vasculature and the density of pericytes in the brain. To clarify whether BDNF acts directly on pericytes, we co-stained BDNF receptor TrkB and the *Pdgfrβ* promoter-driven reporter tdTomato in the brain tissue. TrkB was expressed in a subset of pericytes, as shown by the colocalization of green and red fluorescence (Fig. 5, A). TrkB-expressing pericytes appeared to be enriched in the corpus callosum region compared to the cortex (Fig. 5, A). We analyzed the publicly available single-cell sequencing dataset [29]. The TrkB-encoding gene *Ntrk2* was transcribed in a sub-group of pericytes, with transcription levels lower in 18-month-old C57BL/6J mice than in 2-month-old control mice (Fig. 5, B; Mann-Whitney U test, *p* < 0.001). *Ntrk2* transcription in endothelial cells was also reduced in 18-month-old mice compared to 2-month-old mice (Fig. 5, B; Mann-Whitney U test, *p* < 0.001). Notably, the level of *Ntrk2* transcription in pericytes was comparable to that in endothelial cells. Endothelial cells have been reported to express TrkB [17]. The *Ngfr* gene, which encodes the p75 neurotrophin receptor, a receptor with low affinity for BDNF, was transcribed at a very low level in either pericytes or endothelial cells (Fig. 5, C).

**Figure 5.**
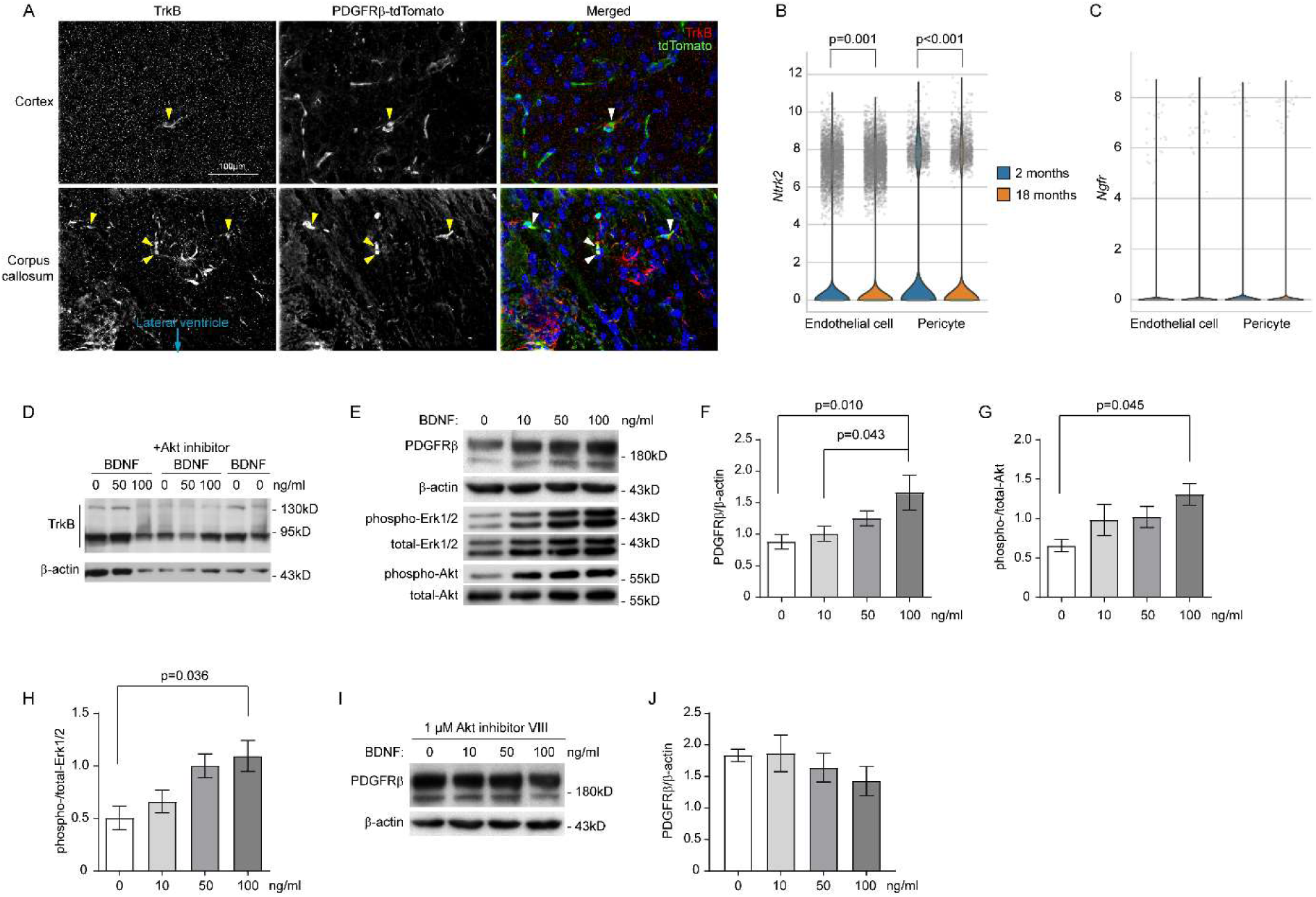
BDNF acts directly on pericytes. Brain sections from 9-month-old pericyte-tdTomato reporter mice were stained for TrkB and tdTomato. TrkB colocalizes with or surrounds tdTomato, particularly at corpus callosum region (A). Single-cell sequencing experiments also showed that the transcription of *Ntrk2* gene decreases in both pericytes and endothelial cells of 18-month-old C57BL/6J mice compared to 2-month-old control mice (B; Mann-Whitney U test, *n* = 5,813 and 11,374 pericytes, 22,898 and 28,556 endothelial cells, for 2- and 18-month-old mice, respectively). The *Ngfr* gene was transcribed at a very low level in both pericytes and endothelial cells (C; Mann-Whitney U test, *p* > 0.05). In following experiments, human pericyte cell lines were cultured. Western blot showed expression of TrkB in the pericyte cell line (D). Cell were then treated with BDNF at 0, 10, 50, and 100 ng/ml for 24 h. Western blot was used to detect the protein level of PDGFRβ, which shows that BDNF treatments significantly increase PDGFRβ expression (E and F; One-way ANOVA followed by Bonferroni *post hoc* test, *n* = 10 per group for 0, 10 and 50 concentrations and *n* = 5 for 100 ng/ml concentration. Ten experiments were independently repeated), as well as to determine the phosphorylation levels of both Akt and Erk1/2 (E, G and H; One-way ANOVA followed by Tukey *post hoc* test, *n* = 4 or 3 per group. Four and three experiments were independently repeated for Akt and Erk1/2, respectively). Finally, pericytes were treated with BDNF at different concentrations in the presence or absence of 1µM Akt inhibitor VIII. The protein level of PDGFRβ as detected by Western blot was not altered by the treatments of BDNF (I and J; One-way ANOVA, *p* > 0.05, *n* = 4 per group. Four experiments were independently repeated).

In additional experiments (see Supplementary Figure 2), pericytes from both young and aged mice were pooled to obtain sufficient cell numbers for reliable analysis. These pericytes were then classified into two groups with (*Ntrk2*_high) and without (*Ntrk2*_zero) transcription of the *Ntrk2* gene. The *Ntrk2*_high group accounted for approximately 15% of all pericytes (2,664 out of 17,187). Compared with the *Ntrk2*_zero group, the *Ntrk2*_high group exhibited 281 DEGs with significant upregulation (log₂ fold change > 1, adjusted *p* < 0.05), whereas no DEGs were found to be significantly downregulated (log₂ fold change < -1, adjusted *p* < 0.05) (see Supplementary Table 1). Gene Ontology and KEGG enrichment analyses revealed that many upregulated DEGs were neuron-specific and associated with synaptic function. Likewise, the transcriptional level of *Aqp4*, an astrocyte-specific gene, was 3.5-fold higher in the high-expression group compared to the low-expression group. Previous studies have shown that AQP4 is more strongly expressed in astrocytic endfoot membranes adjacent to pericytes than in those facing endothelial cells [35]. Together, these findings suggest that TrkB-expressing pericytes are positioned in closer proximity to neurons and astrocytes than TrkB-negative pericytes, as indicated by the neuronal and astrocytic transcripts captured in the single-cell sequencing of pericytes.

We cultured the human pericyte cell line HBPC/ci37 as previously described [27], and observed the expression of TrkB by Western blot (Fig. 5, D). It should be noted that the truncated forms of TrkB were more pronounced than full-length TrkB. Truncated TrkB typically lacks tyrosine kinase activity and inhibits the function of full-length TrkB by competing for ligands via the homologous extracellular domain [36]. Interestingly, we still observed that BDNF increased PDGFRβ expression in a dose-dependent manner after treating cells with recombinant BDNF at concentrations of 0, 10, 50, and 100 ng/mL for 24 hours (Fig. 5, E and F; one-way ANOVA, *p* < 0.05). We also found that BDNF increased the phosphorylation of Akt and Erk1/2 in a concentration-dependent pattern (Fig. 5, E, G and H; one-way ANOVA, *p* < 0.05).

In the following experiment, we investigated whether activation of Akt mediated the effect of BDNF on the expression of PDGFRβ. Cultured pericytes were treated with BDNF at different concentrations with and without the presence of 1 μM Akt inhibitor VIII for 24 hours. The inhibitor at this concentration has been shown in our previous study to inhibit Akt signaling [27]. As shown in Fig. 5, I and J, treatment of Akt inhibitor abolished BDNF-induced expression of PDGFRβ (one-way ANOVA, *p* > 0.05).

## Discussion

Defects in microcirculation are common pathologies in the aging brain that promote age-related neurodegeneration, i.e., AD [2, 10]. However, the molecular mechanisms that underlie vascular dysfunction during aging, as well as potential strategies for prevention, remain poorly understood. BDNF, a key regulator of neuronal survival and synaptic plasticity, is known to decline with age. Our findings extend this knowledge by suggesting that reduced BDNF signaling may also contribute to pericyte loss and impaired microcirculation in the aging brain. Supporting this conclusion, we demonstrated that (i) both vascular density and expression of pericyte marker proteins PDGFRβ and CD13 in cerebral microvessels decreased in parallel with reduced BDNF activity in aged brains, (ii) transient neuronal or astrocytic BDNF deletion for three months was sufficient to reduce vasculature and pericyte numbers in 10-month-old mice, and (iii) pericytes expressed BDNF receptor TrkB, which decreased during aging, and recombinant BDNF directly enhanced PDGFRβ expression in cultured pericytes through an Akt-dependent mechanism.

To date, there have been no publications directly linking BDNF and microcirculation/pericytes in the brain. However, transcription of the *BDNF* gene decreases in the brain tissue of patients with severe depression [37], which correlates with reduced cerebral blood flow (CBF) particularly in the anterior cingulate and prefrontal cortex [38–40]. It is noteworthy that the reduction in CBF correlates more strongly with trait depression than with state depression, suggesting that reduced CBF is related to a structural predisposition rather than a state-dependent functional change [41]. Outside of the brain, BDNF is expressed throughout the cardiovascular system, including endothelial cells and vascular smooth muscle, and promotes angiogenesis and enhances the capillary density during cardiovascular development [17, 18]. Conditional knockout of TrkB in pericytes/smooth muscle cells (SMCs) under the control of *Smc22α* promoter reduces the coverage of pericytes/SMCs, alters the ultrastructure of endothelial cells and increases vascular permeability in postnatal mice [42]. A prospective study in a large community showed that higher levels of BDNF are associated with a lower risk of cardiovascular events and death, independent of low-grade inflammation, body mass index, physical activity and depression [43]. Our study may have for the first time demonstrated the direct action of BDNF on pericytes, which subsequently affects microvascular circulation in the brain.

It cannot be ruled out that BDNF also regulates the structure and function of the vascular system by protecting endothelial cells [17, 18]. The binding of platelet-derived growth factor (PDGF)-B released by endothelial cells to PDGFRβ on pericytes is essential for the proliferation and integration of pericytes into the blood vessel [44]. We have observed that pericytes and endothelial cells express the BDNF receptor TrkB at comparable levels (see Fig. 5, A and B). It should be noted that only a subset of pericytes or endothelial cells express TrkB. Mechanisms that drive the development of the TrkB-positive cell population and maintain this population need to be investigated. Interestingly, single-cell sequencing analyses revealed that TrkB-positive pericytes exhibit a higher abundance of neuron- and astrocyte-specific transcripts compared to TrkB-negative pericytes. This discrepancy is unlikely to be attributed to technical artifacts, as both groups of cells should have an equal likelihood of transcript contamination. These findings raise the possibility that TrkB-enriched pericytes reside in closer proximity to neurons and astrocytes - for example, within the neurovascular unit - thereby rendering them more prone to incorporating neuronal and astrocytic RNA from the surrounding microenvironment. Nevertheless, the potential spatial and functional relationship between TrkB-expressing pericytes and neurons/astrocytes requires further validation using complementary approaches, such as electron microscopy.

We have observed that the Akt signaling pathway mediates the effects of BDNF in pericytes. The protein content of mature BDNF always correlates positively with Akt phosphorylation and PDGFRβ and CD13 expression in both the brain and cultured pericytes, which is consistent with previous studies [16]. Our previous experiments showed that inhibition of Akt reduces the expression of PDGFRβ and CD13 in cultured pericytes in a dose-dependent manner [27]. Our current study indicated that inhibition of Akt abolishes the effect of BDNF on PDGFRβ expression in pericytes. In neurons, BDNF activates Erk1/2 [45]. We observed activation of Erk1/2 signaling in association with BDNF in the brain and in cultured pericytes; however, this appears to be less significant than the effect of BDNF on Akt. For example, Erk1/2 phosphorylation in the brain was not reduced upon neuronal BDNF deficiency and even increased in the aged brain. Since the activation of Akt and Erk1/2 generally promotes cell survival and proliferation, we can assume that BDNF is capable of maintaining the homeostasis of pericytes as well as the structure and function of the microvascular network in the aged brain.

How brain aging regulates BDNF-TrkB signaling in pericytes is unclear. It is known that inflammatory activation increases in the aging brain (e.g. through upregulation of *Il-1β* and *Tnf-α* gene transcription) [46]. In cultured cortical neurons, withdrawal of growth factors such as B-27 or fetal calf serum leads to cell death, which can be prevented by BDNF. Cotreatment with IL-1β abolishes this protective effect of BDNF, possibly by impairing the binding of signaling molecules, i.e., PI3K/Akt, to TrkB [47]. Our previous studies have shown that haploinsufficiency of MyD88 in microglia reduces neuroinflammation and improves microvasculature in the brains of APP/PS1 transgenic mice [26]. Thus, it is worth investigating whether neuroinflammation regulates BDNF-TrkB signaling in pericytes.

Regular physical activity is a well-known strategy for maintaining brain health. Physical activity increases BDNF levels in the blood and has a positive effect on the structure, function and cognitive abilities of the brain in older adults [48]. In 5×FAD mice, exercise elevates BDNF expression in the hippocampus. Overexpression of BDNF in combination with adult neurogenesis in the hippocampus mimics the effects of exercise on improving cognitive function [49]. Interestingly, aerobic exercise is also associated with an increase in cerebral blood flow [50–52]. A recent meta-analysis of six population-based cohort studies with a total of 8517 participants showed that physical activity is associated with lower WMH volume, larger total brain volume and a lower risk of dementia [53]. Similarly, aerobic exercise can also increase red blood cell flow and oxygen supply in the capillaries of the subcortical region of mice [54]. Therefore, our study could suggest that the action of BDNF on pericytes represents a link between exercise and microvascular circulation in the brain.

## Conclusion

Our findings demonstrate that diminished BDNF signaling - whether driven by aging or neuronal and astrocytic *Bdnf* gene knockout - leads to a reduction in pericyte density and cerebral microvasculature. Conversely, enhancing BDNF expression, either pharmacologically or through regular physical exercise, holds promise for restoring pericyte function and improving microcirculatory dynamics in the aging brain. These results advance our understanding of the vascular components of brain aging and highlight BDNF as a potential therapeutic target for preventing vascular decline in older adults and possibly in aging-related disorders such as AD.

## Data availability

All data generated or analyzed during this study are included in this published article and available from the corresponding author on reasonable request.

## List of abbreviations

AD: Alzheimer’s disease
BDNF: Brain-derived neurotrophic factor
BBB: Blood-brain barrier
CreERT2: Cre recombinase and an estrogen receptor ligand binding domain
CSF: Cerebrospinal fluid
DEG: Differentially expressed gene
HBSS: Hanks’ balanced salt solution
HBPC: Human brain vascular pericyte
KEGG: Kyoto Encyclopedia of Genes and Genomes
PDGFRβ: Platelet-derived growth factor receptor
β PBS: Phosphate-buffered saline
RIPA: Radioimmunoprecipitation assay buffer
SMC: Smooth muscle cell
TrkB: Tyrosine kinase B receptor
WMH: White matter hyperintensities

## Acknowledgments

We thank Dr. M. Sendtner (University of Würzburg) for providing *Bdnf*-floxed mice, and Dr. F. Kirchhoff (Saarland University) for *Gfap*-CreERT2 transgenic mice. We appreciate Elisabeth Gluding and Kati Jordan for their excellent technical assistance.

## Author contributions

Y.L. conceptualized and designed the study, acquired funding, conducted experiments, acquired and analyzed data, and wrote the manuscript. Q.L., W.Q., Q.C., W.H., C. S. and Z.T. conducted experiments, acquired data and analyzed data. T.F. provided cell lines. G.P. and Z.T. discussed the study concept and designed the project. All authors contributed to the article and approved the submitted version.

## Ethics declarations

Ethics approval and consent to participate

Animal experiments were conducted in accordance with national rules and ARRIVE guidelines, and authorized by Landesamt für Verbraucherschutz, Saarland, Germany (registration numbers: 06/2017 and 05/2022).

## Consent for publication

Not applicable.

## References

1. Kelly-Hayes M: Influence of age and health behaviors on stroke risk: lessons from longitudinal studies. J Am Geriatr Soc 2010, 58 Suppl 2(Suppl 2):S325–328. doi:10.1111/j.1532-5415.2010.02915.x

2. Wardlaw JM, Smith EE, Biessels GJ, Cordonnier C, Fazekas F, Frayne R, Lindley RI, O’Brien JT, Barkhof F, Benavente OR et al: Neuroimaging standards for research into small vessel disease and its contribution to ageing and neurodegeneration. Lancet Neurol 2013, 12(8):822–838. doi:10.1016/S1474-4422(13)70124-8

3. Debette S, Markus HS: The clinical importance of white matter hyperintensities on brain magnetic resonance imaging: systematic review and meta-analysis. BMJ 2010, 341:c3666. doi:10.1136/bmj.c3666

4. Kloppenborg RP, Nederkoorn PJ, Geerlings MI, van den Berg E: Presence and progression of white matter hyperintensities and cognition: a meta-analysis. Neurology 2014, 82(23):2127–2138. doi:10.1212/WNL.0000000000000505

5. Fang M, Hu J, Weiss J, Knopman DS, Albert M, Windham BG, Walker KA, Sharrett AR, Gottesman RF, Lutsey PL et al: Lifetime risk and projected burden of dementia. Nat Med 2025, 31(3):772–776. doi:10.1038/s41591-024-03340-9

6. Maillard P, Fletcher E, Carmichael O, Schwarz C, Seiler S, DeCarli C, Alzheimer’s Disease Neuroimaging I: Cerebrovascular markers of WMH and infarcts in ADNI: A historical perspective and future directions. Alzheimers Dement 2024, 20(12):8953–8968. doi:10.1002/alz.14358

7. Strain JF, Phuah CL, Adeyemo B, Cheng K, Womack KB, McCarthy J, Goyal M, Chen Y, Sotiras A, An H et al: White matter hyperintensity longitudinal morphometric analysis in association with Alzheimer disease. Alzheimers Dement 2023, 19(10):4488–4497. doi:10.1002/alz.13377

8. Wardlaw JM, Smith C, Dichgans M: Small vessel disease: mechanisms and clinical implications. Lancet Neurol 2019, 18(7):684–696. doi:10.1016/S1474-4422(19)30079-1

9. Dupre N, Drieu A, Joutel A: Pathophysiology of cerebral small vessel disease: a journey through recent discoveries. J Clin Invest 2024, 134(10):e172841. doi:10.1172/JCI172841

10. Bennett HC, Zhang Q, Wu YT, Manjila SB, Chon U, Shin D, Vanselow DJ, Pi HJ, Drew PJ, Kim Y: Aging drives cerebrovascular network remodeling and functional changes in the mouse brain. Nat Commun 2024, 15(1):6398. doi:10.1038/s41467-024-50559-8

11. Decker Y, Muller A, Nemeth E, Schulz-Schaeffer WJ, Fatar M, Menger MD, Liu Y, Fassbender K: Analysis of the vasculature by immunohistochemistry in paraffin-embedded brains. Brain Struct Funct 2018, 223(2):1001–1015. doi:10.1007/s00429-017-1595-8

12. Sweeney MD, Ayyadurai S, Zlokovic BV: Pericytes of the neurovascular unit: key functions and signaling pathways. Nat Neurosci 2016, 19(6):771–783. doi:10.1038/nn.4288

13. Cicognola C, Mattsson-Carlgren N, van Westen D, Zetterberg H, Blennow K, Palmqvist S, Ahmadi K, Strandberg O, Stomrud E, Janelidze S et al: Associations of CSF PDGFRbeta With Aging, Blood-Brain Barrier Damage, Neuroinflammation, and Alzheimer Disease Pathologic Changes. Neurology 2023, 101(1):e30–e39. doi:10.1212/WNL.0000000000207358

14. Nation DA, Sweeney MD, Montagne A, Sagare AP, D’Orazio LM, Pachicano M, Sepehrband F, Nelson AR, Buennagel DP, Harrington MG et al: Blood-brain barrier breakdown is an early biomarker of human cognitive dysfunction. Nat Med 2019, 25(2):270–276. doi:10.1038/s41591-018-0297-y

15. Faraji J, GA SM: Harnessing BDNF Signaling to Promote Resilience in Aging. Aging Dis 2024, 16(4):1813–1841. doi:10.14336/AD.2024.0961

16. Colucci-D’Amato L, Speranza L, Volpicelli F: Neurotrophic Factor BDNF, Physiological Functions and Therapeutic Potential in Depression, Neurodegeneration and Brain Cancer. Int J Mol Sci 2020, 21(20):7777. doi:10.3390/ijms21207777

17. Donovan MJ, Lin MI, Wiegn P, Ringstedt T, Kraemer R, Hahn R, Wang S, Ibanez CF, Rafii S, Hempstead BL: Brain derived neurotrophic factor is an endothelial cell survival factor required for intramyocardial vessel stabilization. Development 2000, 127(21):4531–4540. doi:10.1242/dev.127.21.4531

18. Zierold S, Buschmann K, Gachkar S, Bochenek ML, Velmeden D, Hobohm L, Vahl CF, Schafer K: Brain-Derived Neurotrophic Factor Expression and Signaling in Different Perivascular Adipose Tissue Depots of Patients With Coronary Artery Disease. J Am Heart Assoc 2021, 10(6):e018322. doi:10.1161/JAHA.120.018322

19. Lautrup S, Myrup Holst C, Yde A, Asmussen S, Thinggaard V, Larsen K, Laursen LS, Richner M, Vægter CB, Prieto GA et al: The role of aging and brain-derived neurotrophic factor signaling in expression of base excision repair genes in the human brain. Aging Cell 2023, 22(9):e13905. doi:10.1111/acel.13905

20. Rauskolb S, Zagrebelsky M, Dreznjak A, Deogracias R, Matsumoto T, Wiese S, Erne B, Sendtner M, Schaeren-Wiemers N, Korte M et al: Global deprivation of brain-derived neurotrophic factor in the CNS reveals an area-specific requirement for dendritic growth. J Neurosci 2010, 30(5):1739–1749. doi:10.1523/JNEUROSCI.5100-09.2010

21. Madisen L, Zwingman TA, Sunkin SM, Oh SW, Zariwala HA, Gu H, Ng LL, Palmiter RD, Hawrylycz MJ, Jones AR et al: A robust and high-throughput Cre reporting and characterization system for the whole mouse brain. Nat Neurosci 2010, 13(1):133–140. doi:10.1038/nn.2467

22. Hirrlinger PG, Scheller A, Braun C, Hirrlinger J, Kirchhoff F: Temporal control of gene recombination in astrocytes by transgenic expression of the tamoxifen-inducible DNA recombinase variant CreERT2. Glia 2006, 54(1):11–20. doi:10.1002/glia.20342

23. Hao W, Luo Q, Menger MD, Fassbender K, Liu Y: Treatment With CD52 Antibody Protects Neurons in Experimental Autoimmune Encephalomyelitis Mice During the Recovering Phase. Front Immunol 2021, 12:792465. doi:10.3389/fimmu.2021.792465

24. Cuervo H, Pereira B, Nadeem T, Lin M, Lee F, Kitajewski J, Lin C-S: PDGFRβ-P2A-CreERT2 mice: a genetic tool to target pericytes in angiogenesis. Angiogenesis 2017, 20(4):655–662. doi:10.1007/s10456-017-9570-9

25. Hao W, Luo Q, Tomic I, Quan W, Hartmann T, Menger MD, Fassbender K, Liu Y: Modulation of Alzheimer’s disease brain pathology in mice by gut bacterial depletion: the role of IL-17a. Gut Microbes 2024, 16(1):2363014. doi:10.1080/19490976.2024.2363014

26. Quan W, Luo Q, Hao W, Tomic I, Furihata T, Schulz-Schaffer W, Menger MD, Fassbender K, Liu Y: Haploinsufficiency of microglial MyD88 ameliorates Alzheimer’s pathology and vascular disorders in APP/PS1-transgenic mice. Glia 2021, 69(8):1987–2005. doi:10.1002/glia.24007

27. Quan W, Luo Q, Tang Q, Furihata T, Li D, Fassbender K, Liu Y: NLRP3 Is Involved in the Maintenance of Cerebral Pericytes. Front Cell Neurosci 2020, 14:276. doi:10.3389/fncel.2020.00276

28. Zudaire E, Gambardella L, Kurcz C, Vermeren S: A computational tool for quantitative analysis of vascular networks. PLoS One 2011, 6(11):e27385. doi:10.1371/journal.pone.0027385

29. Jin K, Yao Z, van Velthoven CTJ, Kaplan ES, Glattfelder K, Barlow ST, Boyer G, Carey D, Casper T, Chakka AB et al: Brain-wide cell-type-specific transcriptomic signatures of healthy ageing in mice. Nature 2025, 638(8049):182-196. doi:10.1038/s41586-024-08350-8

30. Luecken MD, Buttner M, Chaichoompu K, Danese A, Interlandi M, Mueller MF, Strobl DC, Zappia L, Dugas M, Colome-Tatche M et al: Benchmarking atlas-level data integration in single-cell genomics. Nat Methods 2022, 19(1):41–50. doi:10.1038/s41592-021-01336-8

31. Wu T, Hu E, Xu S, Chen M, Guo P, Dai Z, Feng T, Zhou L, Tang W, Zhan L et al: clusterProfiler 4.0: A universal enrichment tool for interpreting omics data. Innovation (Camb) 2021, 2(3):100141. doi:10.1016/j.xinn.2021.100141

32. Umehara K, Sun Y, Hiura S, Hamada K, Itoh M, Kitamura K, Oshima M, Iwama A, Saito K, Anzai N et al: A New Conditionally Immortalized Human Fetal Brain Pericyte Cell Line: Establishment and Functional Characterization as a Promising Tool for Human Brain Pericyte Studies. Molecular Neurobiology 2018, 55(7):5993–6006. doi:10.1007/s12035-017-0815-9

33. Ahmad N, Wang W, Nair R, Kapila S: Relaxin induces matrix-metalloproteinases-9 and -13 via RXFP1: induction of MMP-9 involves the PI3K, ERK, Akt and PKC-zeta pathways. Mol Cell Endocrinol 2012, 363(1-2):46–61. doi:10.1016/j.mce.2012.07.006

34. Jelinic M, Marshall SA, Leo CH, Parry LJ, Tare M: From pregnancy to cardiovascular disease: Lessons from relaxin-deficient animals to understand relaxin actions in the vascular system. Microcirculation 2019, 26(2):e12464. doi:10.1111/micc.12464

35. Gundersen GA, Vindedal GF, Skare O, Nagelhus EA: Evidence that pericytes regulate aquaporin-4 polarization in mouse cortical astrocytes. Brain Struct Funct 2014, 219(6):2181–2186. doi:10.1007/s00429-013-0629-0

36. Li Y, Wei C, Wang W, Li Q, Wang ZC: Tropomyosin receptor kinase B (TrkB) signalling: targeted therapy in neurogenic tumours. J Pathol Clin Res 2023, 9(2):89–99. doi:10.1002/cjp2.307

37. Ray MT, Shannon Weickert C, Webster MJ: Decreased BDNF and TrkB mRNA expression in multiple cortical areas of patients with schizophrenia and mood disorders. Transl Psychiatry 2014, 4(5):e389. doi:10.1038/tp.2014.26

38. Li J, Yang Y, Zhu Y, Zhou L, Han Y, Yin T, Cheng Z, Zhang G, Shen Y, Chen J: Towards characterizing the regional cerebral perfusion in evaluating the severity of major depression disorder with SPECT/CT. BMC Psychiatry 2018, 18(1):70. doi:10.1186/s12888-018-1654-6

39. Monkul ES, Silva LA, Narayana S, Peluso MA, Zamarripa F, Nery FG, Najt P, Li J, Lancaster JL, Fox PT et al: Abnormal resting state corticolimbic blood flow in depressed unmedicated patients with major depression: a (15)O-H(2)O PET study. Hum Brain Mapp 2012, 33(2):272–279. doi:10.1002/hbm.21212

40. Vasic N, Wolf ND, Gron G, Sosic-Vasic Z, Connemann BJ, Sambataro F, von Strombeck A, Lang D, Otte S, Dudek M et al: Baseline brain perfusion and brain structure in patients with major depression: a multimodal magnetic resonance imaging study. J Psychiatry Neurosci 2015, 40(6):412–421. doi:10.1503/jpn.140246

41. Chiappelli J, Adhikari BM, Kvarta MD, Bruce HA, Goldwaser EL, Ma Y, Chen S, Ament S, Shuldiner AR, Mitchell BD et al: Depression, stress and regional cerebral blood flow. J Cereb Blood Flow Metab 2023, 43(5):791–800. doi:10.1177/0271678X221148979

42. Anastasia A, Deinhardt K, Wang S, Martin L, Nichol D, Irmady K, Trinh J, Parada L, Rafii S, Hempstead BL et al: Trkb signaling in pericytes is required for cardiac microvessel stabilization. PLoS One 2014, 9(1):e87406. doi:10.1371/journal.pone.0087406

43. Kaess BM, Preis SR, Lieb W, Beiser AS, Yang Q, Chen TC, Hengstenberg C, Erdmann J, Schunkert H, Seshadri S et al: Circulating brain-derived neurotrophic factor concentrations and the risk of cardiovascular disease in the community. J Am Heart Assoc 2015, 4(3):e001544. doi:10.1161/JAHA.114.001544

44. Lindahl P, Johansson BR, Leveen P, Betsholtz C: Pericyte loss and microaneurysm formation in PDGF-B-deficient mice. Science 1997, 277(5323):242–245. doi:10.1126/science.277.5323.242

45. Fukunaga K, Miyamoto E: Role of MAP kinase in neurons. Mol Neurobiol 1998, 16(1):79–95. doi:10.1007/BF02740604

46. Letiembre M, Hao W, Liu Y, Walter S, Mihaljevic I, Rivest S, Hartmann T, Fassbender K: Innate immune receptor expression in normal brain aging. Neuroscience 2007, 146(1):248–254. doi:10.1016/j.neuroscience.2007.01.004

47. Tong L, Balazs R, Soiampornkul R, Thangnipon W, Cotman CW: Interleukin-1 beta impairs brain derived neurotrophic factor-induced signal transduction. Neurobiol Aging 2008, 29(9):1380–1393. doi:10.1016/j.neurobiolaging.2007.02.027

48. Walsh EI, Smith L, Northey J, Rattray B, Cherbuin N: Towards an understanding of the physical activity-BDNF-cognition triumvirate: A review of associations and dosage. Ageing Res Rev 2020, 60:101044. doi:10.1016/j.arr.2020.101044

49. Choi SH, Bylykbashi E, Chatila ZK, Lee SW, Pulli B, Clemenson GD, Kim E, Rompala A, Oram MK, Asselin C et al: Combined adult neurogenesis and BDNF mimic exercise effects on cognition in an Alzheimer’s mouse model. Science 2018, 361(6406):eaan8821. doi:10.1126/science.aan8821

50. Chapman SB, Aslan S, Spence JS, Defina LF, Keebler MW, Didehbani N, Lu H: Shorter term aerobic exercise improves brain, cognition, and cardiovascular fitness in aging. Front Aging Neurosci 2013, 5:75. doi:10.3389/fnagi.2013.00075

51. Alfini AJ, Weiss LR, Nielson KA, Verber MD, Smith JC: Resting Cerebral Blood Flow After Exercise Training in Mild Cognitive Impairment. J Alzheimers Dis 2019, 67(2):671–684. doi:10.3233/JAD-180728

52. Smith KJ, Ainslie PN: Regulation of cerebral blood flow and metabolism during exercise. Exp Physiol 2017, 102(11):1356–1371. doi:10.1113/EP086249

53. Grasset L, Bis JC, Frenzel S, Kojis D, Simino J, Yaqub A, Beiser A, Berr C, Bressler J, Bulow R et al: Selected social and lifestyle correlates of brain health markers: the Cross-Cohort Collaboration Consortium. Alzheimers Dement 2025, 21(4):e70148. doi:10.1002/alz.70148

54. Shin P, Pian Q, Ishikawa H, Hamanaka G, Mandeville ET, Guo S, Fu B, Alfadhel M, Allu SR, Sencan-Egilmez I et al: Aerobic exercise reverses aging-induced depth-dependent decline in cerebral microcirculation. Elife 2023, 12:e86329. doi:10.7554/eLife.86329

