## Supplementary Figures for "Brain-derived neurotrophic factor supports pericyte and vascular homeostasis in the aging brain"

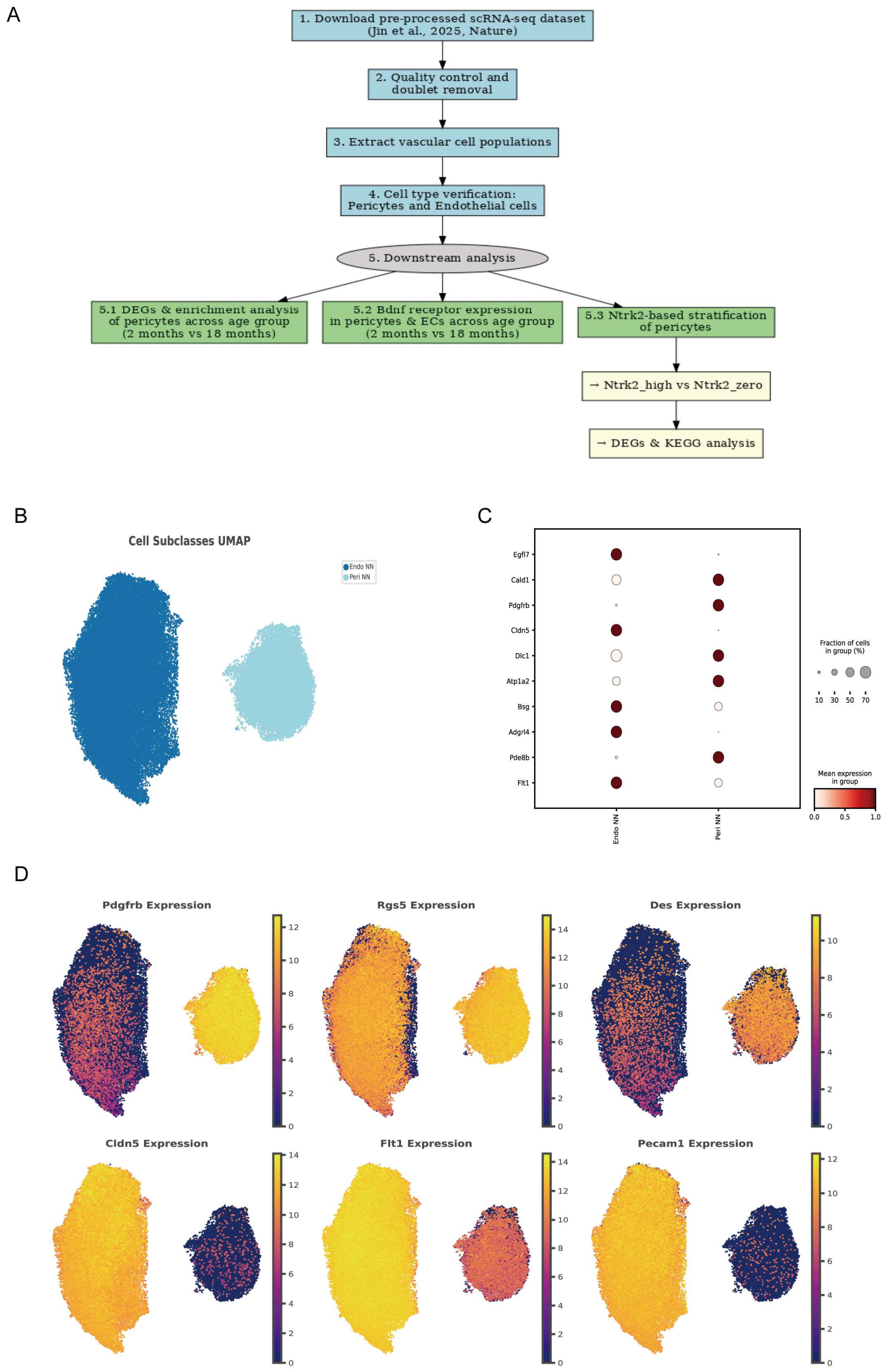


**Supplementary Figure 1, (A)** Workflow of scRNA-seq data analysis. Overview of the analysis pipeline including extraction of vascular cells, identification of pericytes and endothelial cells, age group comparison (2 months vs. 18 months), assessment of BDNF receptor expression, and *Ntrk2*-based stratification with downstream enrichment analysis. **(B)** UMAP visualization of extracted endothelial cells and pericytes, showing clear separation of these two vascular cell types in low-dimensional space. **(C)** Dot plot showing the top five marker genes for each vascular cell type. Dot size represents the percentage of cells expressing the gene, and color intensity reflects the average expression level. **(D)** UMAP feature plots displaying three canonical pericyte markers and three canonical endothelial markers. Yellow indicates high expression (high enrichment), and purple indicates low expression.

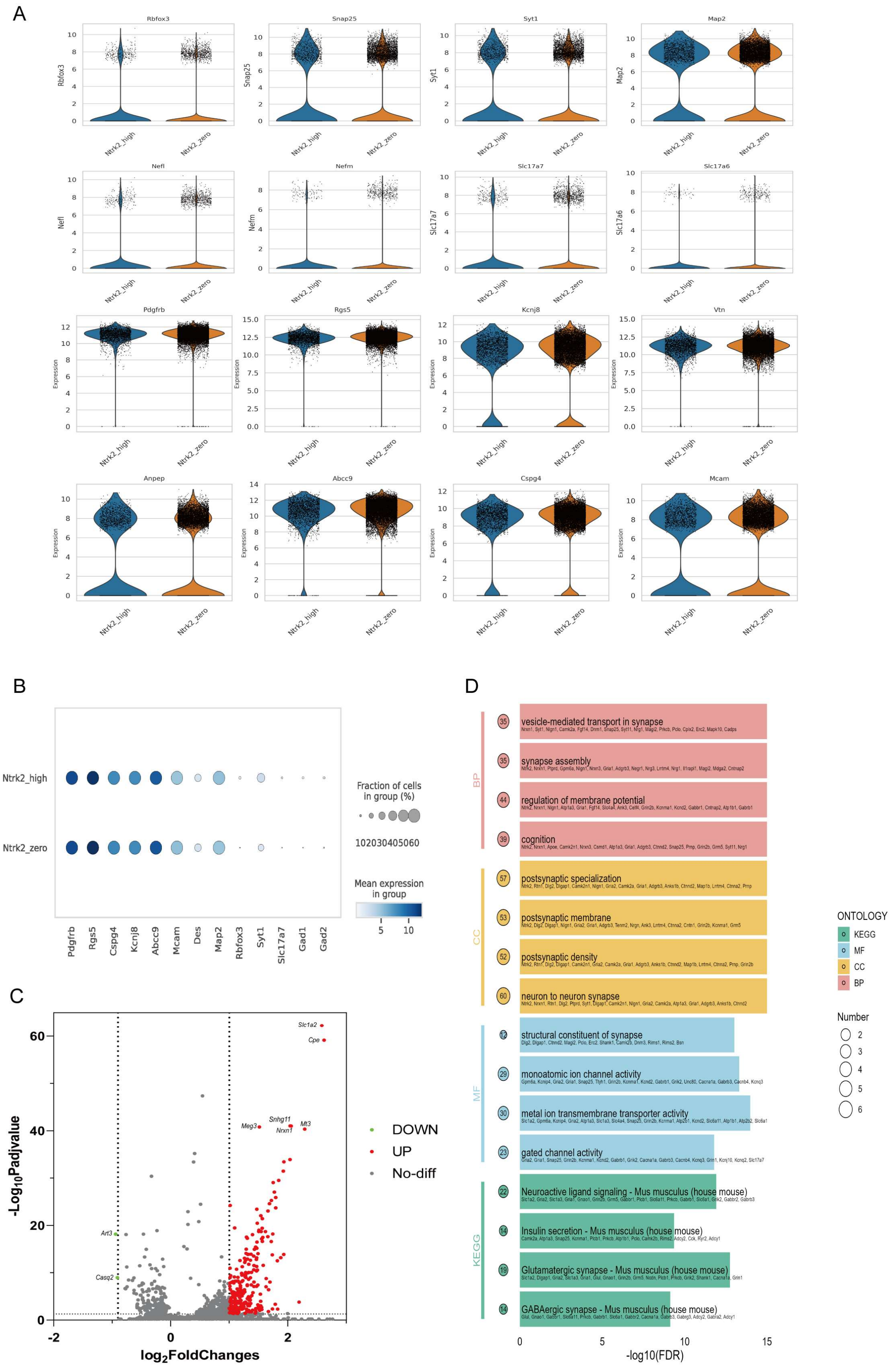
A
Rbfox3
Snap25
Syt1
Map2
Nefl
Nefm
Slc17a7
Slc17a6
Pdgfrb
Rgs5
Kcnj8
Vtn
Anpep
Abcc9
Cspg4
Mcam
Expression
Ntrk2_high
Ntrk2_zero
B
Fraction of cells
in group (%)
102030405060
Mean expression
in group
5
10
Pdgfrb
Rgs5
Cspg4
Kcnj8
Abcc9
Mcam
Des
Map2
Rbfox3
Syt1
Slc17a7
Gad1
Gad2
C
Slc1a2
Cpe
Meg3
Snhg11
Nrxn1
Mt3
Art3
Casq2
DOWN
UP
No-diff
-Log10Padjvalue
log2FoldChanges
D
BP
vesicle-mediated transport in synapse
synapse assembly
regulation of membrane potential
cognition
CC
postsynaptic specialization
postsynaptic membrane
postsynaptic density
neuron to neuron synapse
MF
structural constituent of synapse
monoatomic ion channel activity
metal ion transmembrane transporter activity
gated channel activity
KEGG
Neuroactive ligand signaling - Mus musculus (house mouse)
Insulin secretion - Mus musculus (house mouse)
Glutamatergic synapse - Mus musculus (house mouse)
GABAergic synapse - Mus musculus (house mouse)
-log10(FDR)
ONTOLOGY
KEGG
MF
CC
BP
Number

**Supplementary Figure 2, (A)** Violin plots of marker gene expression stratified by *Ntrk2*-defined groups. Cells were divided into "*Ntrk2*_high" (*Ntrk2* > 0) and "*Ntrk2*_zero" (*Ntrk2* = 0). The plotted genes include general neuronal markers (e.g., *Rbfox3*, *Snap25*, *Syt1*, *Map2*, *Nefl*, *Nefm*, *Slc17a7*, *Slc17a6*) and general pericyte markers (e.g., *Pdgfrb*, *Rgs5*, *Kcnj8*, *Vtn*, *Anpep*, *Abcc9*, *Cspg4*, *Mcam*). Single-cell values are shown. Plots were generated using Scanpy's violin function with jittered strip plots overlaid to show individual-cell distributions. **(B)** Dot plot showing pericyte and neuronal marker expression across "*Ntrk2*_high" and "*Ntrk2*_zero" groups. Pericyte markers included *Pdgfrb*, *Rgs5*, *Cspg4*, *Kcnj8*, *Abcc9*, *Mcam*, and *Des*; neuronal markers included *Map2*, *Rbfox3*, *Syt1*, *Slc17a7*, *Gad1*, and *Gad2*. Color intensity represents average expression after normalization ($\log_2$(CPM + 1)), and dot size indicates the fraction of cells expressing the gene within each group. Generated with Scanpy's dotplot. **(C)** Volcano plot of differential gene expression in pericytes (*Ntrk2*_high vs *Ntrk2*_zero). The x-axis shows $\log_2$ fold change ($\log_2$FC) and the y-axis shows $-\log_{10}$(FDR). Significance was defined using asymmetric effect-size thresholds with FDR control: up-regulated genes have $\log_2FC \geq 1$ and FDR < 0.05 (red); down-regulated genes have $\log_2FC \leq -0.9$ and FDR < 0.05 (green); others are non-significant (gray). Dashed lines indicate thresholds (vertical: $x = 1$ and $x = -0.9$; horizontal: $y = -\log_{10}(0.05) = 1.301$). Representative genes (e.g., *Slc1a2*, *Cpe*) are labeled. *Ntrk2* itself shows an extremely large $\log_2$FC and lies outside the plotting range; it is therefore not displayed. **(D)** KEGG/GO enrichment dot plots in pericytes (*Ntrk2*_high vs *Ntrk2*_-zero). Differential genes were defined as $|\log_2FC| \geq 1$ with FDR < 0.05 and subjected to KEGG and GO enrichment. The x-axis shows $-\log_{10}$(FDR), the y-axis lists enriched terms, and bubble size encodes the gene ratio. Colors indicate the source ontology/database: red = GO Biological Process (BP), yellow = GO Cellular Component (CC), blue = GO Molecular Function (MF), green = KEGG. The top four terms per ontology/database by significance are shown (16 total), summarizing functional shifts and significance patterns for this contrast.
